# Novel Immunoassay for Diagnosis of Ongoing *Clostridioides difficile* Infections Using Serum and Medium Enriched for Newly Synthesized Antibodies (MENSA)

**DOI:** 10.1101/2020.04.23.058859

**Authors:** Natalie S. Haddad, Sophia Nozick, Geena Kim, Shant Ohanian, Colleen Kraft, Paulina A. Rebolledo, Yun Wang, Hao Wu, Adam Bressler, Sang Nguyet Thi Le, Merin Kuruvilla, L. Edward Cannon, F. Eun-Hyung Lee, John L. Daiss

## Abstract

**BACKGROUND:** *Clostridioides difficile* infections (CDI) have been a challenging and increasing serious concern in recent years. While early and accurate diagnosis is crucial, available assays have frustrating limitations

**OBJECTIVE:** Develop a simple, blood-based immunoassay to accurately diagnose patients suffering from active CDI.

**MATERIALS AND METHODS:** Uninfected controls (n=95) and CDI patients (n=167) were recruited from Atlanta area hospitals. Blood samples were collected from patients within twelve days of a positive CDI test and processed to yield serum and PBMCs cultured to yield medium enriched for newly synthesized antibodies (MENSA). Multiplex immunoassays measured Ig responses to ten recombinant *C. difficile* antigens.

**RESULTS:** Sixty-six percent of CDI patients produced measurable responses to *C. difficile* antigens in their serum or MENSA within twelve days of a positive CDI test. Fifty-two of the 167 CDI patients (31%) were detectable in both serum and MENSA, but 32/167 (19%) were detectable only in MENSA, and 27/167 (16%) were detectable only in serum.

**DISCUSSION:** We describe the results of a multiplex immunoassay for the diagnosis of ongoing CDI in hospitalized patients. Our assay resolved patients into four categories: MENSA-positive only, serum-positive only, MENSA- and serum-positive, and MENSA- and serum-negative. The MENSA positive-only patients accounted for 30% and may be attributed to nascent antibody secretion in MENSA prior to seroconversion. Conversely, the serum positive-only subset may have been more advanced in their disease course. Immunocompromise and misdiagnosis may have contributed to the 34% of CDI patients who were not identified using MENSA or serum immunoassays.

**IMPORTANCE:** While there was considerable overlap between patients identified through MENSA and serum, both methods detected additional, unique patients. The combined use of both MENSA and serum to detect CDI patients resulted in the greatest identification of CDI patients. Together, longitudinal analysis of MENSA and serum will provide a more accurate evaluation of successful host humoral immune responses in CDI patients.

## INTRODUCTION

*Clostridioides difficile* infections (CDI) afflict nearly half a million patients in the United States on an annual basis, and have progressively increased in incidence as well as severity in recent years. These infections lead to the death of 29,000 patients in the United States annually, most of whom are elderly (1, 2). While typically regarded a hospital acquired entity, the epidemiology of CDI has shifted in the past decade and half the cases are now community acquired among low risk populations (3). Another particular challenge in recent years is the high rate of recurrent CDI, up to 20-35% within eight weeks of the primary infection (4). This risk increases up to 65% following secondary and tertiary infections (5). Due to the high incidence of primary infection and the frequency of recurrence, CD represents a substantial healthcare burden with annual costs estimated to exceed five billion dollars annually in the United States alone (6, 7).

Current methods of diagnosis test stool samples for the presence of secretory proteins such as glutamate dehydrogenase (GDH) or toxin TcdB, or the CD bacteria itself by measuring characteristic genes, e.g., TcdB. However, the diagnosis of CDI in hospitals is confounded by multiple factors. First, five percent of the general population and more than twenty percent of the hospitalized population are asymptomatic carriers (8-11). Second, even among hospitalized patients, diarrhea is caused by CDI in only 10-20% of cases, despite concurrent leukocytosis or immune compromise (12, 13). Third, the CD bacterium itself may be shifting in response to treatment strategies. Frequently distinguished by the secretion of the toxin TcdA only twenty years ago (14), CD pathogens now frequently secrete the other signature toxin, TcdB. Further, a third multi-subunit toxin, binary toxin (CDT), has become more common in recent years (15, 16). No single currently available diagnostic test has acceptable specificity. To circumvent these limitations, many hospitals use a multi-tier diagnostic strategy that typically includes initial screening for the presence of secretory products in the stool followed by PCR for characteristic CD genes. However, the sole reliance on molecular diagnostics without querying the host response will likely result in false positive diagnoses and unwarranted overtreatment. Recent literature underscores the potential for overtreatment despite this multi-step approach (17).

The development of specific serum antibodies in response to ongoing CDI is well recognized (14, 18). However, the utility of measuring the antibody response is confounded by the sub-population of patients who carry serum antibodies consequent to environmental exposure and subclinical infection (2, 19, 20). Further, the increase in serum anti-CD antibody levels may occur well into the infectious course, typically in the second week following symptom onset (21). Finally, titers also persist long after resolution of the underlying infection and are thus of limited specificity.

Recently, we and others have described the rapid and potent generation of newly stimulated antibody secreting cells (ASC) in the circulation during the window between infection onset and prior to seroconversion for bacterial and viral pathogens (22-25). These ASCs produce large quantities of pathogen-specific antibodies, and their detection may provide a broadly applicable approach to analyze the host humoral immune response during an acute infection.

In contrast to serum anti-CD antibodies, the ASC response is transient and coincides with the acute infectious phase. Newly stimulated in the lymph nodes, ASC are typically absent prior to infection. They travel to lymphoid organs such as the bone marrow via the circulation and disappear following symptom resolution (22). Preparation of ASC from an anti-coagulated blood sample is straightforward. One simply: 1) harvests the ASC-containing peripheral blood mononuclear cells (PBMC) from the blood; 2) washes away contaminating Ig from the serum; and 3) cultures the washed PBMC to allow them to secrete their antibodies into the culture medium creating a novel sample we call medium enriched for newly synthesized antibodies (MENSA).

In this observational investigation, we present the results of a pilot study validating the use of a multi-antigen immunoassay and its ability to measure antibody secretion reflecting ongoing CDI in MENSA samples from a complex hospital population and compare its performance with results from sera.

## RESULTS

### Selection, synthesis, and characterization of candidate *C. difficile* antigens

Ten CD antigens were selected primarily by reported immunogenicity and secretion or surface expression (Table 1). The secreted toxins were prepared as selected constituent domains (e.g., the glucosyltransferase domain (GTD) of TcdA) or as subunits (e.g., CDTa or CDTb). Cwp84 (26-29) and flagellin (30) were each chosen based on their potential utility as vaccine targets. GDH was selected because of its widespread use as a biomarker of CDI (31-33). Each selected antigen was prepared as a His_6_-tagged or GST fusion recombinant protein, expressed in *E. coli*, and purified by chelation chromatography. Following SDS-PAGE, each antigen appeared predominantly as a single band of the anticipated molecular weight (Fig. 1). In addition, available monoclonal antibodies specific for TcdB-CROP/TcdBvir-CROP (34), TcdA-CROP (34), CDTa (35), CDTb (35) and GDH (Invitrogen MA5-18193) reacted exclusively with their corresponding recombinant antigens (Fig. 2 A-C). For the remaining antigens (Cwp84, TcdB-GTD, TcdA-GTD, and flagellin), pooled positive sera responded two orders of magnitude higher compared to pooled negative sera (Fig. 2D). Together, these data demonstrate that each antigen was pure, the correct size, and immunocompetent.

**Table 1.** Molecular Weight, Functions, and Immunogenicity of Ten *C. difficile* Antigens.

| Antigen | MW (kDa) |  | Function | Immunogenicity |
| --- | --- | --- | --- | --- |
|  | Original | Recombinant |  |  |
| TcdA-CROP | 99 | 102 | Subunits of Toxin A (TcdA) and Toxin B (TcdB) are most extensively studied and accepted as major virulence factors in *C. difficile* literature. Cause extensive damage to large intestine by disrupting actin cytoskeleton and impairing tight junctions in intestinal epithelial cells via proinflammatory and cytotoxic activity (59). Combined repetitive oligopeptides (CROP) of each toxin are binding domains that reside on the C-termini of TcdA and TcdB (60). TcdBvir-CROP is the TcdB-CROP region of virulent strain (53, 61) | TcdA-CROP Mice immunized with high amounts of TcdA-CROP monoclonal antibodies protected against challenge with toxigenic strains of *C. difficile* (62). |
| TcdB-CROP | 61 | 64 |  |  |
| TcdBvir-CROP | 61 | 64 |  |  |
| TcdA GTD | 63 | 66 | N-terminal glucosyltransferase (GTD), binds to and inactivates epithelial intracellular Rho GTPases by glucosylation, disrupts cytoskeleton, causes necrosis of the colonic monolayer (52) | TcdB-GTD effective in protecting both cells in invitro cytotoxicity assay and mice upon lethal *C. difficile* toxin challenge, when combined with anti-TcdA-CROP (63). Coadministration of TcdA-CROP & TdB GTD protected animals from *C. difficile* challenge by inducing systemic neutralizing IgG (64). |
| TcdB GTD | 63 | 66 |  |  |
| CDTa | 53 | 49 | Subunits of Binary Toxin AKA *C. difficile* transferase (CDT): Third toxin produced by hypervirulent strains associated with increased severity of CDI; induces depolymerization of actin cytoskeleton, produces microtubule-based membrane protrusions forming network on epithelial cells, increasing bacterial adherence (65, 66). CDTa is the enzymatic ADP-ribosyltransferase, modifies actin. CDTb binds host cells and translocates CDTa into the cytosol. | Mice challenged with native TcdA, TcdB, and binary toxin, immunization with inactive versions all three toxins required for protection. CDTa only partially protected against binary toxin, whereas CDTb alone or with CDTa afforded full protection (67). Protection against CDTa and CDTb required for challenge against BI17/NAP1/BI/027. |
| CDTb | 99 | 121 |  |  |
| GDH | 46 | 47 | Glutamate Dehydrogenase (GDH): NAD-specific enzyme converts L-glutamate into α-ketoglutarate; highly conserved among *C. difficile* ribotypes; often initial test for *C. difficile* in stool (31, 32); highly sensitive but not specific for toxin-producing *C. difficile* isolates; part of a two or three step diagnostic algorithm (68). | No direct evidence for immunogenicity yet. To be determined by us. |
| Cwp84 | 87 | 85 | Secreted cysteine protease; cleaves *C. difficile* paracrystalline surface layer proteins (SLP); cleaves host structural proteins fibronectin, vitronectin and laminin (26-28, 69). | Immunized hamsters 26% weaker & slower colonization and 33% greater survival (28). FliC, FliD, Cwp84 & Cwp66 antibodies found in *C. difficile* patient sera (70). |
| FliC | 31 | 34 | Flagella, composed of flagellin (FliC) and a flagellar cap protein (FliD), enables chemotaxis & adherence to epithelial cell surface (motility & colonization); involved in biofilm formation and toxin production; highly conserved across *C. difficile* strains (20, 55, 56, 71-76) | Immunization of FliC or anti-FliC polyclonal serum protective in mice (30). FliC, FliD, Cwp84 & Cwp66 antibodies found in *C. difficile* patient sera (70) |

**Figure 1.**
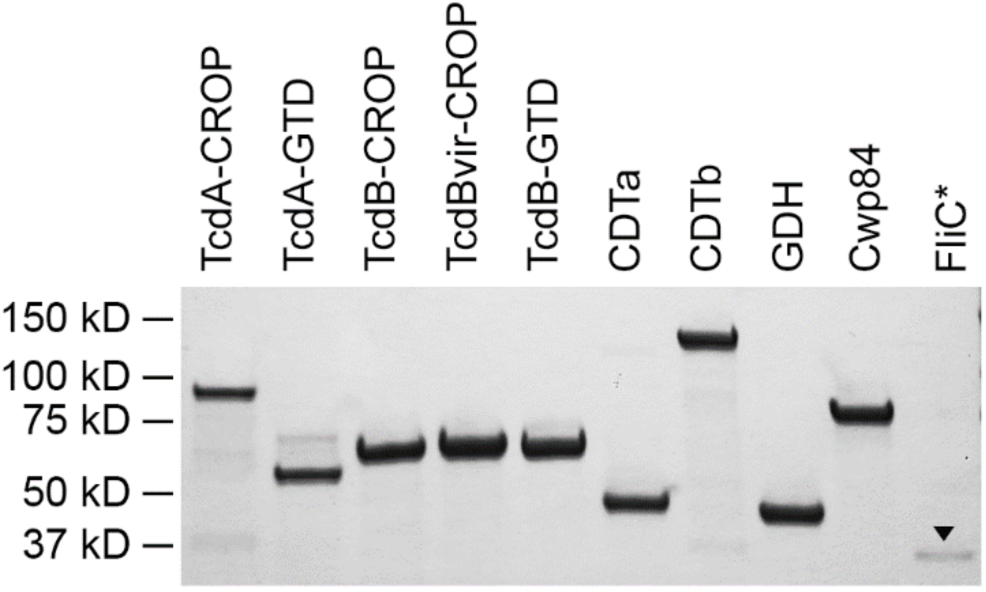
*C. difficile* antigens are pure and display the expected molecular weights. SDS-PAGE confirmed expected molecular weights for recombinant antigens. Each antigen (2.5 µL of 0.5 mg/mL solution) was applied to its assigned well and detected using Bio-Safe Coomassie stain following electrophoresis. All antigens have similar band intensities denoting equal concentrations, except for FliC that started at a much lower concentration (0.1 mg/mL) and yielded a faint band at the anticipated molecular weight (designated with black arrowhead).

**Figure 2.**
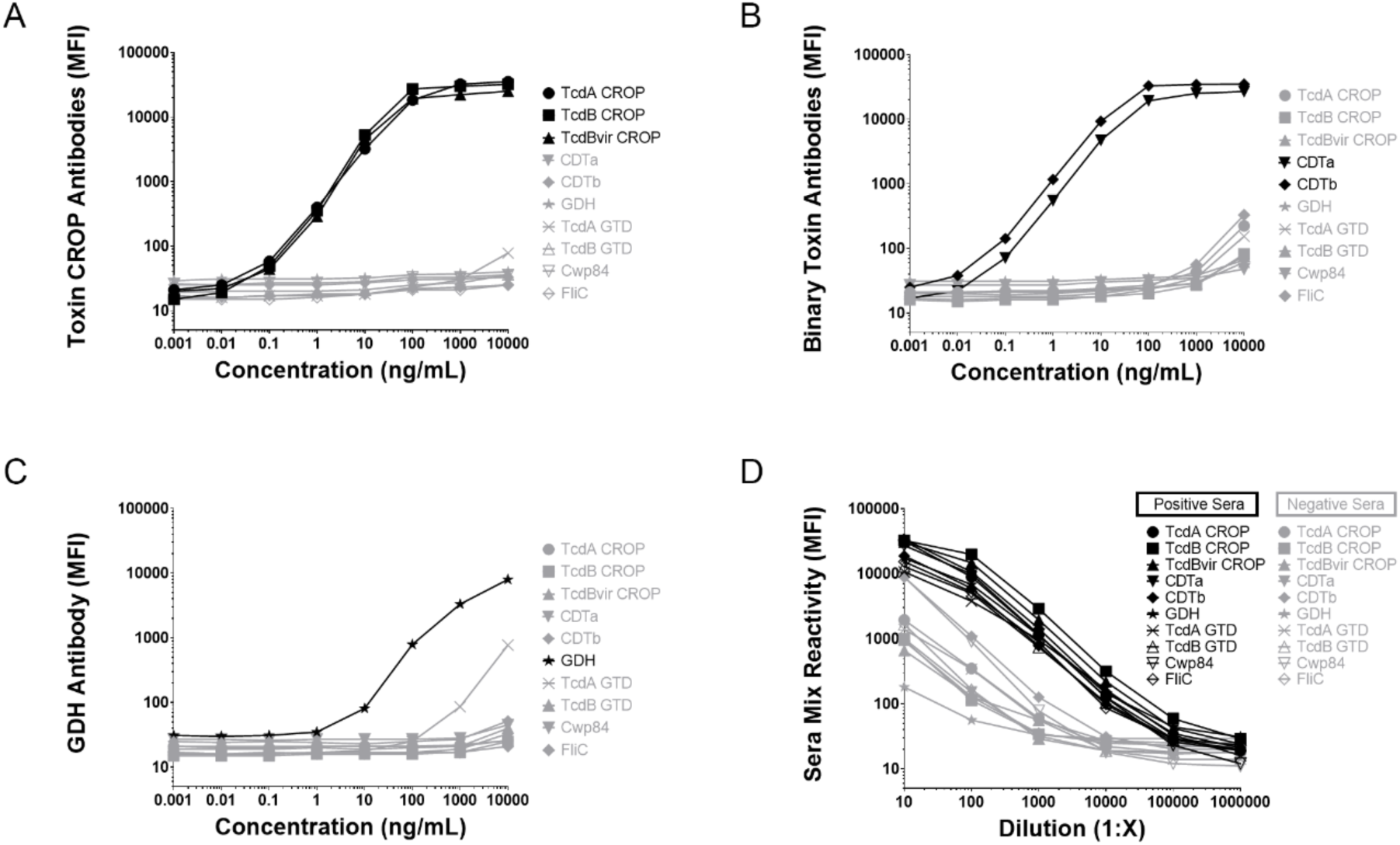
Recombinant CD antigens interact specifically with established monoclonal antibodies and a mixture of positive sera. The 10 CD antigens were tested in MAGPIX-based immunoassays against monoclonal antibodies and/or positive sera. (A) MK-3415 and MK-6072, humanized monoclonal antibodies directed against TcdA-CROP and TcdB-CROP, respectively, were able to specifically detect recombinant antigens between 0.01 ng/mL to 100 ng/mL for TcdA-CROP, TcdB-CROP, and TcdBvir-CROP, with little or no interaction with the other antigens. (B) A mixture of anti-CDTa and anti-CDTb mouse monoclonal antibodies reacted with recombinant CDTa and CDTb in a similar range of detection (0.01 ng/mL to 100 ng/mL) with little or no cross-reactivity with the other antigens. (C) An anti-GDH monoclonal antibody reacted with recombinant GDH between 1-10,000 ng/mL with modest cross-reactivity for TcdA-GTD at higher concentrations. (D) A mixture of *C. difficile* positive patient sera (n=6) shows a range of immuno-reactivity detection for all 10 antigens between dilutions of 1:1,000 to 1:1,000,000. A-C) MFI is a measure of antigen-specific IgG binding to antigens immobilized on designated Luminex beads. D: MFI is the measure of serum-borne antigen-specific IgA+IgG+IgM reacting with each antigen. Sera from healthy donors showed lower reactivity. A-C) Dark lines denote antigen reactivity against their respective monoclonal antibody solution for each panel. D.) Dark lines denote *C. difficile* patient serum reactions against all ten recombinant *C. difficile* antigens, while grey lines denote the much lower reactivity of from healthy donor sera against the 10 *C. difficile* antigens.

### Selection of control subjects and CDI patient characteristics

CDI patients were recruited from Atlanta area hospitals, using the following inclusion criteria. Control subjects were recruited in three groups: 1) healthy donors (H) who had no known exposure to patients with CDI; 2) medical workers (M) who regularly care for CDI patients; and 3) symptomatic patients (S) who had diarrhea but were PCR negative for the TcdB gene in their stool. Serum and MENSA were prepared from each subject and analyzed for antibodies specific for each of the 10 CD antigens using our multiplex immunoassay. Serum and MENSA samples were collected from CDI patients within 12 days of a positive CD test (days post validation; DPV) and were included to illustrate the magnitude of response in each group.

Our final cohort consisted of 167 CDI patients and 95 controls (H=44, M=27, S =24). Demographics of controls were similar to CDI patients in terms of race (*p=0*.*32*) and sex (*p=0*.*51*); however, controls were significantly younger (*p*<*0*.*0001*) (Table 2).

**Table 2.** Demographics of Control and *C. difficile*-infected Subjects

|  | Age |  | Race** |  | Sex |  |
| --- | --- | --- | --- | --- | --- | --- |
|  | < 50 | ≥ 50 | Black | White | Female | Male |
| All Controls | n (%) | n (%) | n (%) | n (%) | n (%) | n (%) |
| Healthy (n=44) | 37 (84) | 7 (16) | 24 (55) | 10 (23) | 30 (68) | 14 (32) |
| Medical (n=27) | 23 (85) | 4 (15) | 20 (74) | 6 (22) | 22 (81) | 5 (19) |
| Symptomatic (n=24) | 12 (50) | 12 (50) | 12 (50) | 9 (38) | 9 (38) | 15 (63) |
| Total (n=95) | 72 (76) | 23 (24) | 56 (59) | 25 (26) | 61 (64) | 34 (36) |
| C<sub>o</sub> Controls |  |  |  |  |  |  |
| Healthy (n=38) | 31 (82) | 7 (18) | 19 (50) | 10 (26) | 26 (68) | 12 (32) |
| Medical (n=26) | 22 (85) | 4 (15) | 19 (73) | 6 (23) | 21 (81) | 5 (19) |
| Total (n=64) | 53 (83) | 11 (17) | 38 (59) | 16 (25) | 47 (73) | 17 (27) |
| C. difficile Patients |  |  |  |  |  |  |
| Pretrial (n=108) | 40 (37) | 68 (63) | 68 (63) | 39 (36) | 64 (59) | 44 (41) |
| Non-recurrent (n=55) | 22 (40) | 33 (60) | 32 (58) | 21 (38) | 33 (60) | 22 (40) |
| Recurrent (n=9) | 2 (22) | 7 (78) | 4 (44) | 5 (56) | 5 (56) | 4 (44) |
| Trial: NR+R (n=64) | 24 (38)* | 40 (63)* | 36 (56) | 26 (41) | 38 (59) | 26 (41) |
| Total (n=172) | 64 (37) | 108 (63) | 104 (60) | 65 (38) | 102 (59) | 70 (41) |
| All Controls vs All CDI patients | $p < 0.0001$ | | $p = 0.26$ | | $p = 0.51$ | |
| Pretrial vs Trial | $p = 1.00$ | | $p = 0.51$ | | $p = 1.00$ | |

\*Percentages add to 101% due to rounding. \*\*Race percentages ≠ 100% due to small sub-population of other races not listed.

### Examination of the control populations

Seventy-eight of 95 (82%) control subjects had no detectable antibody above the C_0_ for any of the 10 antigens in their serum (37/44 healthy; 21/27 medical; 20/24 symptomatic). 87 of 95 (92%) of control subjects had no detectable antibody above the C_0_ for any of the 10 antigens in their MENSA (42/44 healthy; 27/27 medical; 18/24 symptomatic). The pooled positive and negative serum samples used previously (Fig. 2D) elicited responses above and below (respectively) the C_0_ for each antigen at a 1:1,000 dilution.

### Evaluation of CDI patients

Antibodies specific for each of the 10 CD antigens were tested in our multiplex immunoassay for serum and MENSA (Fig. 3). Two essential points are apparent. First, the many CDI patients display substantially higher antibody levels than the controls for each antigen. Second, there is a small minority of controls who have antibody levels overlapping with the CDI patients.

**Figure 3.**
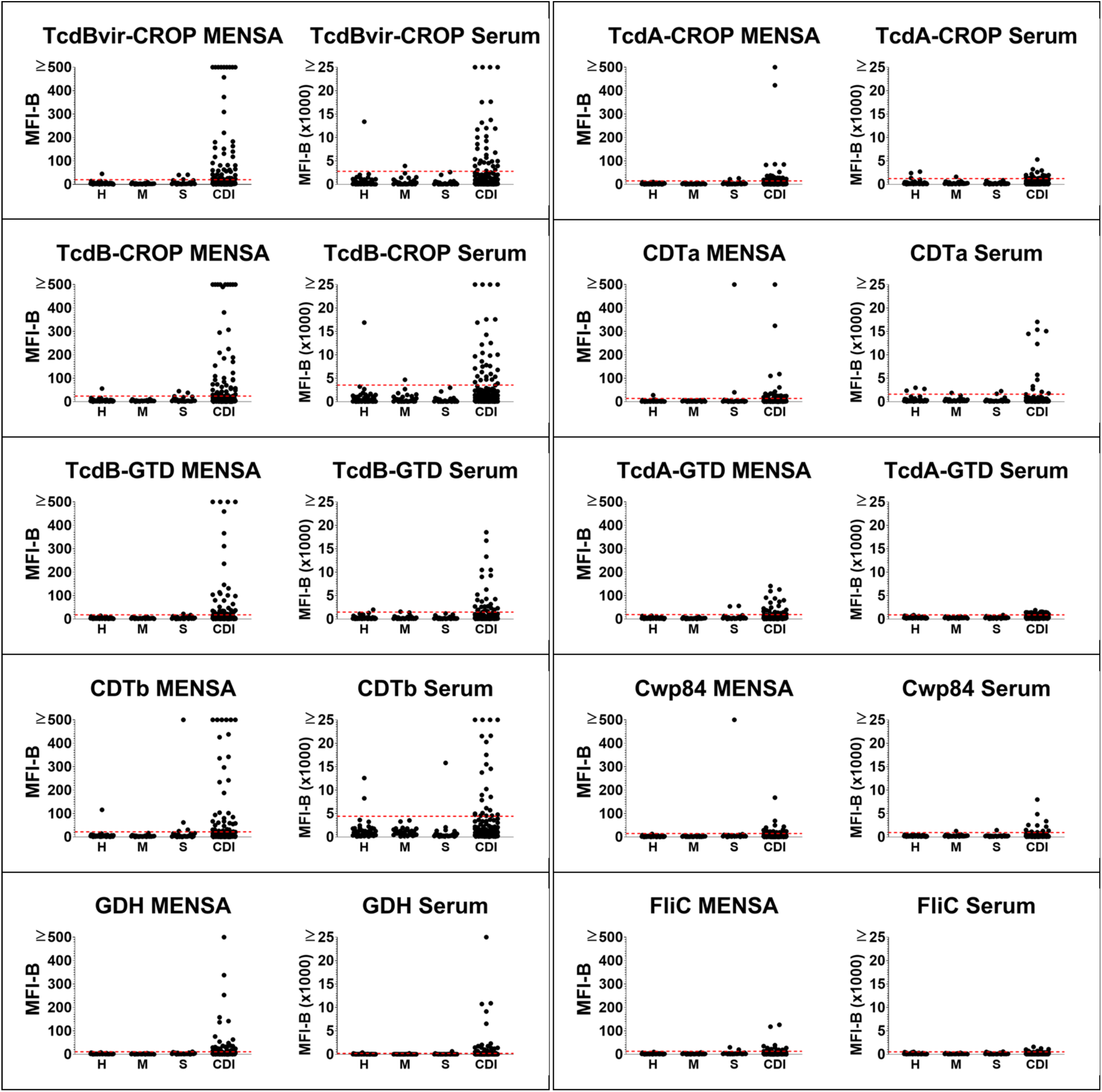
MENSA and Serum responses to 10 *C. difficile* antigens are found at greater frequencies and magnitudes in infected patients compared to control subjects. A) Levels of MENSA antibodies against each of the 10 CD antigens from healthy donors (H; n=44), medical workers (M; n=27), and symptomatic PCR negative patients (S; n=24) compared to CDI patients (CDI; n=167). MFI-B(x1000) value represents the sum of IgA, IgG, and IgM detection. The red dashed line represents the C_0_ value for each antigen.

### Setting C_0_ values

Healthy donor and medical worker controls were used to construct the diagnostic threshold, C_0_. Subjects exhibiting MENSA and/or serum antibody levels greater than ten times the median of the healthy and medical control populations were eliminated from C_0_ determination (denoted by * at the bottom of Fig. 4), leaving 38/44 healthy and 26/27 medical (n=64 total). The C_0_ was calculated for each antigen as the average plus five or four standard deviations of these 64 controls, for MENSA and serum respectively. The symptomatic patients with diarrhea who were PCR negative for TcdB were excluded from the C_0_ calculation due to the possibility that the TcdB PCR test misdiagnoses some CDI cases (e.g., potentially TcdB-negative strains (36, 37)).

**Figure 4.**
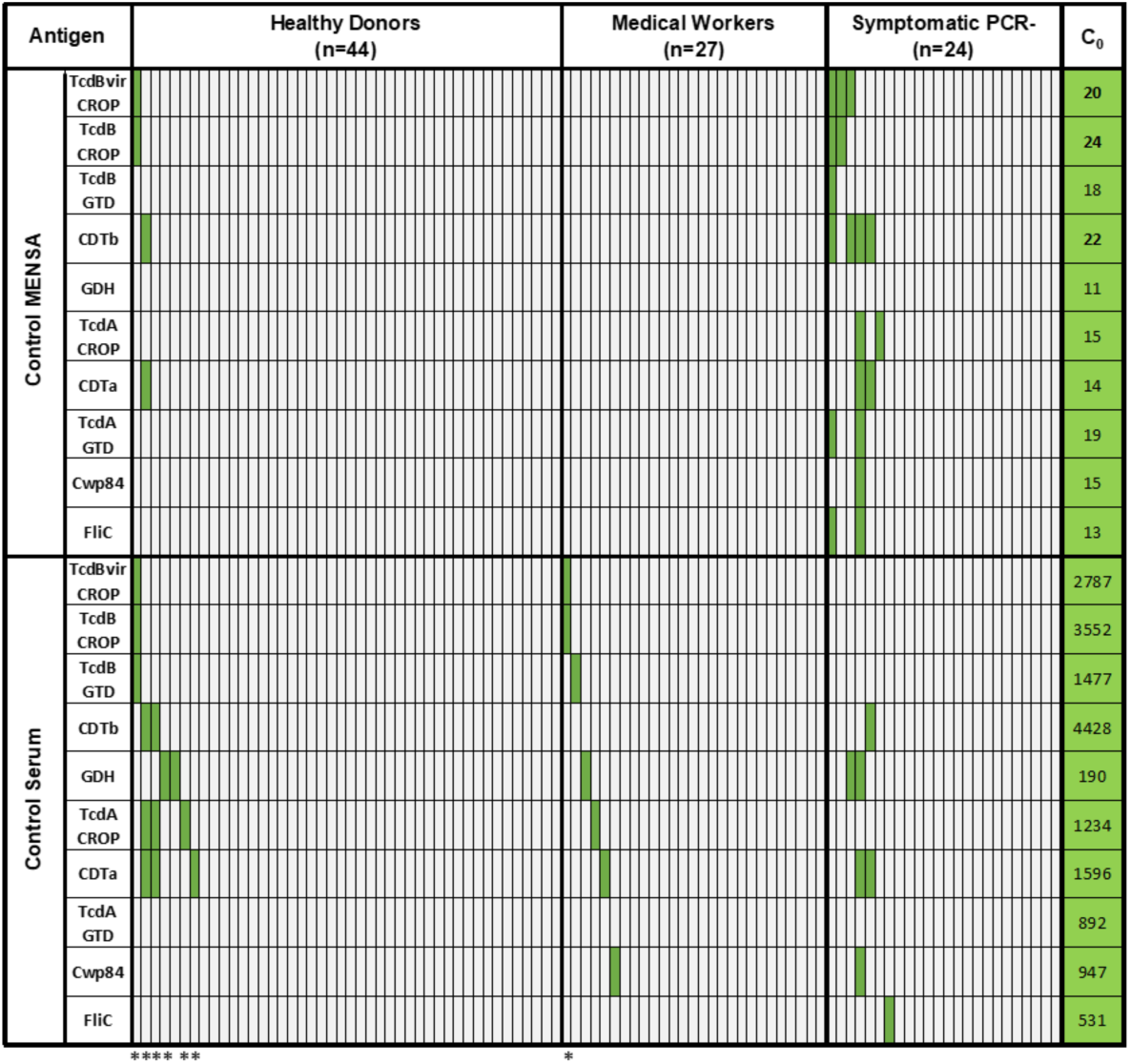
Control subjects negative for *C. difficile* responses are used to construct C_0_ diagnostic thresholds for each antigen. A heat map showing the MENSA (top panel) and serum (bottom panel) responses of all healthy donors, medical workers, and symptomatic PCR negative patients to the 10 CD antigens. Each subject is represented as a single column and each antigen as a single row. The subjects without an asterisk at the bottom (38/44 H and 26/27 M) were used to construct the cutoff values (C_0_): average plus five or four standard deviations in MENSA and serum, respectively. C_0_ values for each antigen are presented in the green column at the right side of the figure. The subjects with an asterisk were eliminated from the C_0_ calculation for having values greater than 10X the median of all healthy and medical subjects. Light grey rectangles represent MFI-B values less than C_0_ for each antigen.

### MENSA and Serum Tests of CDI patients

CDI samples (n=167) selected from patients within 12 DPV responded to CD antigens at higher magnitudes and frequencies than controls (Fig. 3). Eighty-four of 167 (50%) CDI patients exhibited antibody levels above C_0_ for one or more of the 10 CD antigens in their MENSA. Each of the 10 antigens reacted with at least fourteen of the 167 CDI MENSA samples, affirming its immunocompetence. Seventy-nine of 167 (47%) of CDI patients exhibited antibody levels above C_0_ for one or more of the 10 CD antigens in their serum. Positive MENSA and serum samples overlapped in only 31% (52/167) of CDI patients. MENSA alone identified an additional 19% (32/167) of patients, and serum by itself identified an additional 16% (27/167) of patients. MENSA and serum together identified 66% (111/167) of all CDI patients (Fig. 5). Analyzing responses to individual antigens yield smaller subsets of positive CDI patients (7-29%) than a combined ten-antigen test (Fig. 6).

**Figure 5.**
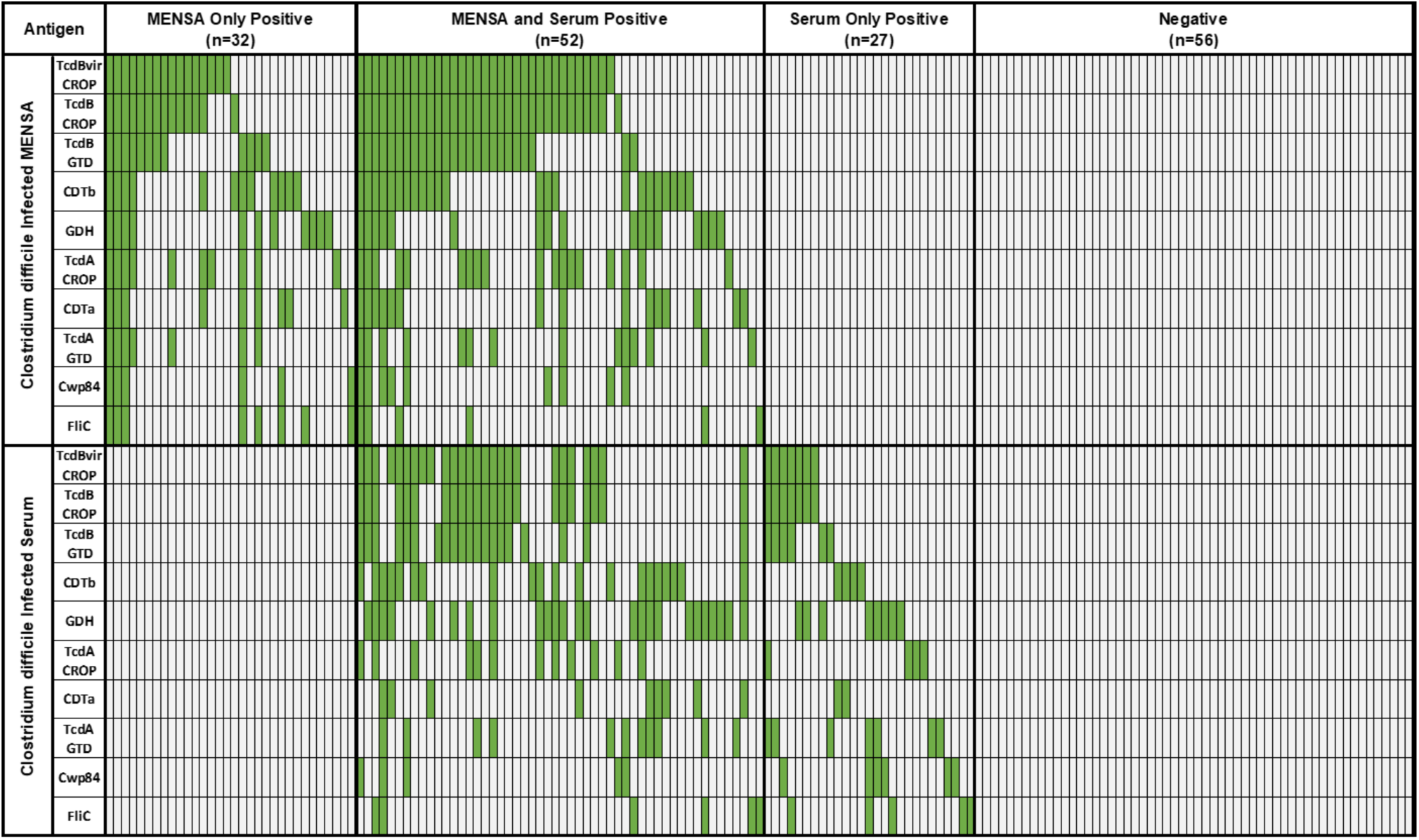
CDI patients can be categorized as positive in MENSA, serum, both, or neither. A heat map showing MENSA and serum responses from 167 CDI patients at a single time point, using the same C_0_ values and color conventions as in Fig. 4. CDI patients are categorized as MENSA Only Positive (n=32), MENSA and Serum Positive (n=52), Serum Only Positive (n=27), or Negative in both sample types (n=56). To facilitate comparison, data for a subject in the MENSA (upper) figure is placed directly above the serum data for the same patient in the serum (lower) figure.

**Figure 6.**
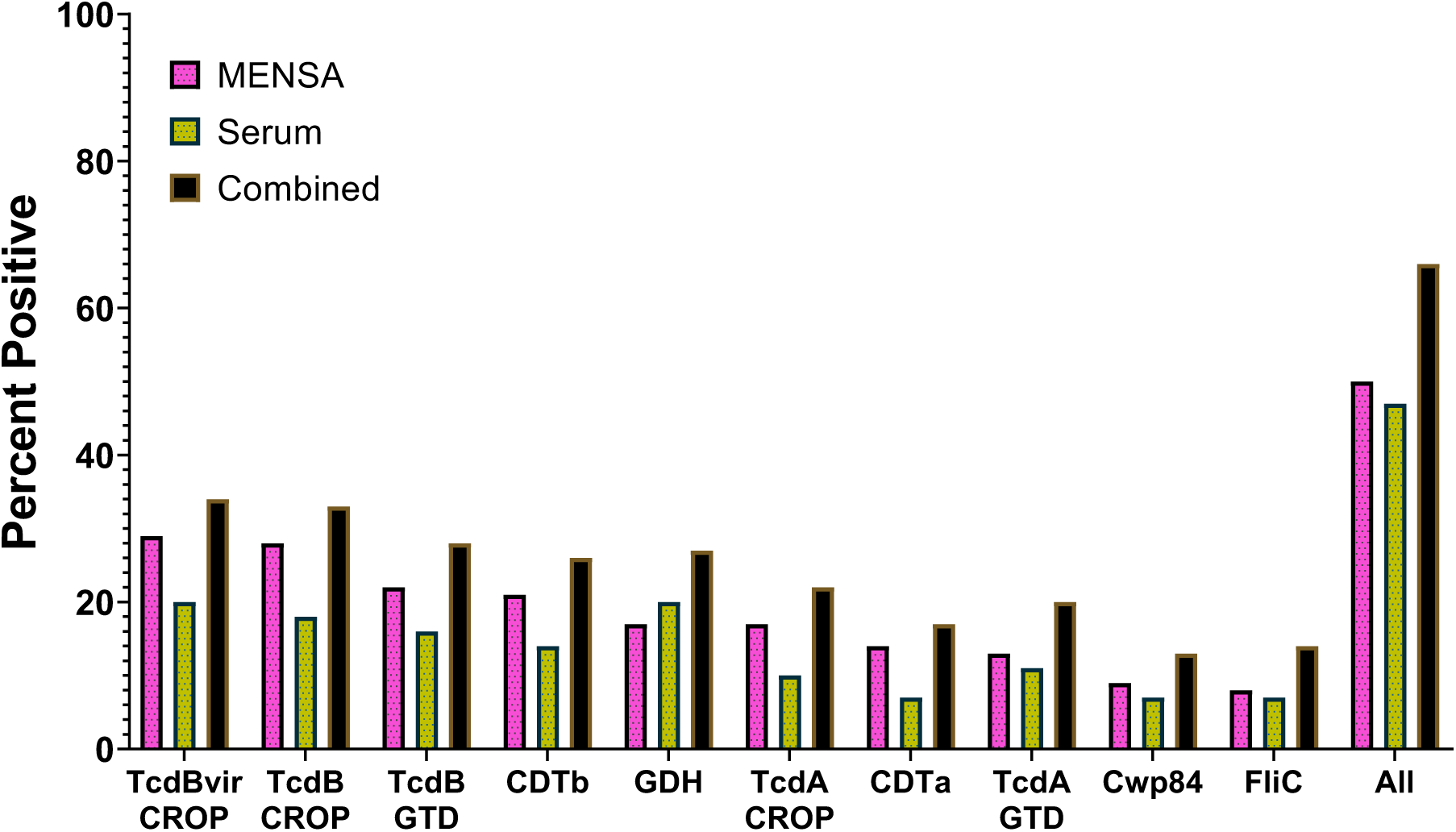
Testing individual antigens in MENSA or serum identifies smaller subsets of CDI patients than testing all antigens together. MENSA and serum samples from infected patients were tested against ten *C. difficile* antigens. Pink bars represent the percentage of CDI patients who were positive for each antigen their MENSA. Yellow bars represent the percentage of CDI patients who were positive for each antigen their serum. Black bars represent the percentage of CDI patients who were positive for each antigen in their MENSA and/or serum.

## DISCUSSION

While the identification of pathogen-specific antibodies is a cornerstone of microbiologic diagnostics, a commercial assay for the serological confirmation of ongoing CDI is lacking. Several investigators have developed tests to evaluate the host humoral immune response to infection (38-40). An association has also been demonstrated between the humoral immune response to CD and infectious course (21). Antibody-based serological testing is hindered by the significant probability of prior environmental exposure to CD and consequent seropositive responses. Further, serum Ig levels can remain elevated for weeks to months post-infection due to the month-long circulating half-life of human IgG and the induction of long-lived plasma cells.

We present the methodology and results of this unique diagnostic approach to detect active CDI, based on measuring the host antibody response in the MENSA and comparing its performance to serum anti-CD antibodies. MENSA allows for the discrimination of newly synthesized ASCs and active Ig secretion, and levels decline rapidly once the inciting infection has resolved. It thus provides a means to circumvent the limitations of serum Ig and affords a high-resolution analysis of the acute antibody response. Our results support the utility of MENSA in acute CDI detection, with a testing format able to achieve high diagnostic specificity in detecting pathogen-specific antibody responses. However, this response is exquisitely time sensitive explaining the significant differences between patient groups and individuals.

### Responses in the control population

Controls were recruited in three groups: healthy, medical, and symptomatic. The symptomatic cohort was not used to establish the C_0_ calculation as they may have included false negatives with CDI undetected by the TcdB PCR test alone. Two of these patients had positive MENSA and serum antibody responses against CDTa and/or CDTb but were negative for TcdB-CROP, suggesting they may have been infected with a binary toxin positive, TcdB negative CD strain (15, 41-43). CD strains positive for binary toxin and negative for TcdB have been recently described and would be missed in our symptomatic cohort since only TcdB PCR testing was performed. Twenty-two percent of medical workers and 16% of healthy donors were positive for CD in their serum antibody population, consistent with the possibility of seroconversion as the result of asymptomatic environmental exposure and/or intestinal colonization (44, 45). In contrast, only two of the healthy controls (4%) and none of the medical workers had positive MENSA antibody responses to CD antigens, demonstrating superior specificity to serum.

### Almost two-thirds of patients had antibody detectable in their MENSA or serum

In samples collected shortly after diagnosis from 167 patients with severe diarrhea and positive testing for CDI, 47% of patients had serum antibody specific for at least one antigen and 32% had antibody specific for two or more antigens. When antibody levels were measured in MENSA, 50% of patients were producing antibody against at least one antigen and 37% against two or more. In total, using serum and MENSA, 111 (67%) out of 167 patients were detectable by their Ig responses. Measurement of the ongoing immune response in CDI patients split the antibody-producing population into three sub-groups: 1) 19% (32 patients) made antibodies only in their MENSA (MENSA+/serum-), 2) 31% (52 patients) had antibody measured in both their MENSA and sera (MENSA+/serum+); and 3) 16% (27 patients) had antibody detected only in their sera (MENSA-/serum+) (Fig. 5). The clinical complexity of our cohort precludes further distinctions among the subgroups. Additionally, there was significant variance in the timing of symptom onset and disease progression prior to CDI diagnosis. We hypothesize that MENSA+/serum-patients had samples drawn earlier in the course of CDI prior to seroconversion. Conversely, the MENSA-/serum+ subset may have been further into their infectious course. An alternative is that these patients may have undergone prior seroconversion and may not be actually experiencing CDI. Further, the MENSA+/serum+ subset may represent those who have mounted a successful immune response and should be unlikely to experience recurrence.

### Patients who were negative for anti-*C. difficile* antibodies in both serum and MENSA

Fifty-six of 167 patients in our population tested negative for anti-CD antibodies in their serum as well as MENSA. Considering the complexity of the Emory patient population among whom many are on immunosuppressive drugs or are otherwise immunocompromised, it is probable that some of the recruited cases had limited capability to mount a measurable antibody response. Furthermore, the primary screening assay for CDI was a PCR measurement of the presence of CD DNA (TcdB) in the stool. Recent reports have challenged the accuracy of PCR testing on a single gene, notably TcdB (12, 46). Concerns have been raised whether the PCR method can distinguish patients with CDI from those who are non-infected carriers. Thus, over-prediction of CDI may have underestimate the evident utility of our analytic approach (47). We anticipate that the exceptional specificity associated with capturing an acute immune response would yield less false positive results, and prevent potential overtreatment in high risk patients. Furthermore, patients enrolled at Emory University Hospitals were tested only for the TcdB gene, and not binary toxin, which may have biased our population towards TcdB responses and missed CDI with CDT-positive strains (15, 41-43).

### Limitations of this study

The goal of the present study was to determine whether antibody production in the serum and/or MENSA can be used to either identify or confirm those patients who are experiencing ongoing CDI. Broadly, we can affirm that the ongoing antibody responses on serum or MENSA are correlated with CDI. That said, there are multiple limitations regarding the recent past history of enrolled patients and the timing of blood draws with respect to the onset of symptoms. The stage of infection with respect to symptom onset must be accurately determined since MENSA has obvious differential outcomes based on timing.

A second limitation is the heterogeneity of our subject population. Some patients were suffering serial recurrences, a sub-population excluded after the first fifty patients had been enrolled. Similarly, in the complex Emory population, some patients were potentially immunocompromised and better control for this was introduced in the second year of recruitment. Also, patients were enrolled from multiple departments and subject to discharge at the discretion of the attending physician. As a result, we were unable to obtain follow-up samples or information from many patients.

Finally, the infecting CD strains were not typed or analyzed. The underlying CD ribotype is known to exert pertinent influence on the host immune response (48). Thus, we have been unable to correlate our immunochemical observations with possible strain-specific humoral responses.

### Study implications

A serologic assay for active CDI is highly desirable due to the limitations of molecular diagnostics and the potential for asymptomatic carriage of toxic strains and even CD toxins(49). CDI diagnosis also becomes especially challenging in immunocompromised populations who frequently have unrelated diarrhea complicated by higher rates of CD colonization (50). A single, highly specific test may help streamline the current multi-tier diagnostic approach and facilitate confirmation in at-risk populations. In view of this, we developed a multiplex platform for the rapid detection of pathogen-specific ASCs and applied it to a novel diagnostic assay for acute CDI.

Beyond its diagnostic value, the detection of CD specific ASCs may have additional practical implications. With the progressive increase in CD incidence and severity, it is of exceeding importance to stratify patients by risk of recurrent or severe disease(51). Deficient antibody responses have been previously associated with recurrent CDI(38). Our assay may thus enable detections of patients at high risk for recurrent CDI, which would allow for timelier, and possibly more effective intervention.

Future studies should expand validation to a larger population with known infecting strains. A more careful longitudinal analysis must also be performed with testing at more frequent intervals to develop a more comprehensive view of the host antibody response.

In conclusion, we have verified that this preliminary design and execution of a diagnostic platform based on antibody detection from ASC in MENSA is a valid technique to identify pathogen-specific antibody responses during acute CDI. Our findings, if reproduced in future studies, may support the monitoring of ASC responses to identify patients with decreased antibody responses who may run the risk of more severe infection.

## MATERIALS AND METHODS

### Enrollment of CDI patients and controls

A total of 167 CDI patients and 95 control subjects were recruited at Emory University and Dekalb Medical Center **(now Emory Decatur Hospital)**. CDI was confirmed by PCR and/or ELISA (C. DIFF QUIK CHEK COMPLETE, Alere North America, Orlando, FL, USA) and blood samples were collected at one-to-three time points (Draw 1 at 1-5 days post-PCR and/or ELISA validation (DPV), Draw 2 at 6-12 DPV, Draw 3 at 21-60 DPV). All patients included in study yielded at least one MENSA and serum sample within 12DPV. A blood sample was obtained from the three populations of control subjects (44 healthy donors, 27 medical workers regularly exposed to CDI, and 26 symptomatic patients who tested PCR negative for *C. difficile*).

### MENSA generation

PBMC were isolated by centrifugation (800xg; 25 minutes) using Lymphocyte Separation Media (Corning). Five washes with RPMI-1640 (Corning) were performed to remove serum immunoglobulins, with erythrocyte lysis (5mL; 5 minutes) after the second wash and cell counting after the fourth. Harvested PBMCs were cultured at 10^6^cells/mL in R10 Media (RPMI-1640, 10% Sigma FBS, 1% Gibco Antibiotic/Antimycotic) on a 12-well sterile tissue culture plate for 24 hours. After incubation, the cell suspension was centrifuged (800xg; 5 minutes) and the supernatant (MENSA) was separated from the PBMC pellet, aliquoted and stored at -80°C.

### Serum generation

Whole blood was collected in 2-4 mL clot activator tubes (BD Vacutainer, Franklin Lakes NJ, USA) and incubated at room temperature for at least 30 minutes. Clot was discarded. Remaining supernatant was centrifuged at 800xg for 10 minutes, removed from pellet, aliquoted and stored at -80°C.

### Selection and preparation of recombinant antigens

Ten CD antigens were selected from literature based on pathogenicity and immunogenicity (Table 1) and sequences were prepared for expression as described below.

CD Toxins A and B (TcdA and TcdB, respectively) are divided into four domains each, the first and last of which were selected for this study: Glucosyltransferase (GTD; amino acids 1-542 of TcdA and 1-543 of TcdB) and the Combined Repetitive Oligopeptides (CROP) binding domain (amino acids 1832– 2710 of TcdA and 1834-2366 of TcdB) (52). These domain sequences were selected from the entire TcdA and TcdB proteins of historical strain VPI 10463 (Accession: P18177.3 and P16154.2, respectively). A hypervirulent epidemic strain, NAP1/B1/027 R20291, varies from the historical strain in the CROP domain sequence (53), and thus an additional virulent CROP antigen, TcdBvir-CROP, was made from this strain as well (1834-2366 Accession: CBE02479.1). A His_6_-tag was placed on the end of each antigen with an Avitag between the sequence and His tag for optional biotinylation; none of the antigens in this study were biotinylated. Tags were positioned on each antigen to replace its missing adjacent domains: tags were placed on N-terminus for CROP domain antigens, and on C-terminus for GTD domains. Final sequences were sent to GenScript for *E. coli* expression (Vector E3) and protein was obtained from supernatant, via one step purification by Ni column.

Binary toxin, also known as *C. difficile* transferase (CDT) is composed of two subunits: CDTa and CDTb. We designed a consensus sequence for CDTa based on five highly conserved sequences from NCBI (>96%; Accession: AEC11560.1, AAB67304.2, AEC11578.1, AEC11575.1, AEC11569.1). The first 43 amino acids, which serve as a signal peptide, were removed and the cysteine at the resulting position 2 was mutated to alanine (1mCDTa) to decrease protein dimerization (35, 54). A His_6_-tag was placed at the C-terminus and the final amino acid sequence was sent to GenScript for *E. coli* expression (pET30a) and purification from cell lysate via one step Ni column. A consensus sequence for CDTb was designed from eight highly conserved sequences (>96%; Accession: AEC11570.1, AEC11585.1, AEC11576.1, AEC11564.1, AEC11558.1, AEC11579.1, AAB67305.1, AEC11561.1). Alignments were carried out using Clustal Omega and consensus sequences were found using Seaview. The first 42 amino acids (signal peptide) were removed to produce Pro-CDTb and expressed as a fusion with glutathione S-transferase (GST) on its N-terminus to facilitate solubility. It has been shown that GST-ProCDTb is recognized by a murine anti-CDTb monoclonal antibody from Merck (35), and thus used as is for our immunoassays. The final protein was expressed in *E. coli* (pGEX-4T-1) and purified from cell lysate.

Flagellin (FliC) has been shown to have cross reactive epitopes from various CD strains (30, 55, 56). The full-length protein sequence for FliC (Accession: AAD46086.1) was obtained from CD strain VPI 10463 (30) with a His_6_-tag and AviTag placed on the N-terminus. The protein was expressed in *E. coli* (E3) and purified from inclusion bodies via one-step purification by Ni column purification.

The CD 630 sequence for cell wall protein Cwp84 (Accession: YP-001089300) was obtained from a patent application for use as a diagnostic marker (57). The first 31 amino acids, which serve as a signal peptide, were removed from the N terminus and a His_6_-tag was placed on the C-terminus. The final protein sequence was expressed in *E. coli* (pET30a) and purified from supernatant of cell lysate via one-step purification by Ni column.

Glutamate dehydrogenase (GDH) sequence (Accession: AAA62756.1) was obtained from Lyerly et al., 1991 (58). A His_6_-tag was placed on the N-terminus, expressed in *E. coli* (pET30a) and purified from supernatant of cell lysate.

### SDS-PAGE

Ten recombinant CD antigens were diluted to 0.5 mg/mL in Phosphate Buffered Saline (Corning Cellgro), or concentrated up to 0.5 mg/mL using 10K MWCO Amicon Ultra-15 centrifugal filters (Millipore), mixed with 2X Laemmli Sample Buffer (Bio-Rad) and 2-mercaptoethanol (Bio-Rad), heated at 100°C for 5 minutes, ran on a 12-well precast 4-15% mini-PROTEAN TGX gel (Bio-Rad), and stained with Bio-Safe Coomassie stain (Bio-Rad), according to BioRad SDS-PAGE protocol.

### Immunoassay validation of antigens

Recombinant antigens were then validated by immunoassay. Human monoclonal antibodies were obtained for TcdA-CROP (MK-3415/GS-CDA1/CDA1/Actoxumab) and TcdB-CROP (MK-6072/MDX-1388/CDB1, Bezlotoxumab; Merck). Mouse monoclonal antibodies against binary toxin subunits CDTa and CDTb and anti-Clostridia GDH monoclonal mouse antibody (Clone: J44D, Invitrogen) were obtained from Merck. A mixture of the human monoclonal antibodies MK-3415 and MK-6072 were titrated and used to test TcdA-CROP, TcdB-CROP, and TcdBvir-CROP antigen immuno-reactivity using antigen-coupled MagPlex® microspheres in a MAGPIX sandwich immunoassay and detected with anti-IgG-PE, as described below. Sequence similarity between the historical and virulent CROP domains of Toxin B allowed MK-6072 to be used for both TcdB-CROP and TcdBvir-CROP. A mixture of the mouse monoclonal anti-CDTa and anti-CDTb was titrated and immuno-reactivities of the CDTa and CDTb antigens were measured. Finally, the mouse monoclonal anti-GDH was titrated and tested against the GDH antigen. Antibodies against the remaining four antigens (TcdA-GTD, TcdB-GTD, FliC, and Cwp84) were unavailable. Therefore, these antigens were tested against a positive serum mixture made from six CD positive patient sera and compared against a healthy donor’s negative serum sample. A MAGPIX sandwich immunoassay was run on the sera mix, with detection by the pooled PE-conjugated (anti-IgA + anti-IgG + anti-IgM).

### C. DIFF QUIK CHEK COMPLETE®

The GDH antigen, Toxin A, and Toxin B were tested against the *C. DIFF QUIK CHEK COMPLETE*® rapid assay (Techlab) according to manufacturer’s protocol and as described in *Alere/Techlab QCC Protocol Overview* Methods section, replacing stool samples with antigen/toxin solutions. The solution was made of 1 ng/mL TcdA-CROP, 1 ng/mL TcdB-CROP, and 2 ng/mL GDH diluted in kit-provided sample diluent.

### Carbodiimide coupling of microspheres to recombinant *C. difficile* antigens

The 10 recombinant CD antigens were separately coupled to microspheres of different regions (Luminex; Austin, TX, USA). Coupling was carried out at room temperature following standard carbodiimide coupling procedures, described below.

Microspheres were washed once with deionized water and incubated for 20 minutes in an end-over-end rotator in the dark in a suspension of 0.1 M NaH_2_PO_4_ (VWR; Radnor, PA, USA), 5 mg/mL EDC (1-Ethyl-3-(3-dimethylaminopropyl) carbodiimide; ThermoFisher Scientific; Waltham, MA, USA), and 5 mg/mL Sulfo-NHS (N-hydroxysulfosuccinimide; ThermoFisher Scientific; Waltham, MA, USA). Activated microspheres were washed twice in 0.05 M MES (2-(N-morpholino)ethanesulfonic acid, Boston Bioproducts, Ashland MA USA) and incubated for two hours in the dark in an end-over-end rotator in a suspension of 0.05 M MES and 1µg/mL antigen. Coupled microspheres were then washed four times in a blocking buffer consisting of 1% BSA (Bovine Serum Albumin; Boston Bioproducts; Ashland, MA, USA), 1X PBS (VWR; Radnor, PA, USA), 0.05 % Sodium azide (VWR; Radnor, PA, USA), and 0.02 % Tween-20 (VWR; Radnor, PA, USA). Coupled microspheres were then stored at a concentration of 10^6^ spheres per 1 mL at 4°C in the same blocking buffer. Concentrations of spheres were confirmed by enumeration with a hemocytometer stage and laboratory microscope at 10X objective.

### Luminex proteomic assays for measurement of anti-antigen antibody

Fifty µL coupled microsphere mix was added to 96-well v-bottom plates (Sorensen BioScience; Murray, UT, USA) at a concentration of 1000 microspheres per region per well. All wash steps and dilutions were accomplished using 1% BSA, 1X PBS assay buffer. MENSA was assayed neat (no dilution) and serum was assayed at 1:1,000 dilution and surveyed for antibodies against recombinant CD antigens. After a one-hour incubation in the dark on a plate shaker at 800 rpm, wells were washed five times in 100 µL of assay buffer, then applied with 3 µg/mL PE-conjugated Goat Anti-Human IgA, IgG and/or IgM (Southern Biotech; Birmingham, AL, USA). After 30 minutes of incubation at 800 rpm in the dark, wells were washed three times in 100 µL of assay buffer, resuspended in 100 µL of assay buffer, and incubated at 800 rpm in the dark for 15 minutes before analysis using a Luminex MAGPIX instrument (Luminex; Austin, TX, USA) running xPonent 4.2 software. Further data manipulation and analysis was undertaken using Microsoft Excel (Microsoft Corporation; Redmond, WA, USA), GraphPad Prism (GraphPad Software; La Jolla, CA, USA), and JMP (SAS Institute Inc.; Cary, NC, USA) statistical analysis packages.

### Data analysis

Median Fluorescent Intensity (MFI) using combined or individual detection antibodies (anti-IgA/anti-IgG/anti-IgM) was measured using the Luminex xPONENT software. The background value of RPMI-160 or assay buffer was subtracted from each MENSA or serum sample result, respectively, to obtain Median Fluorescent Intensity minus Background (MFI-B). Control MENSA and serum samples from healthy donors (n=44) and medical workers (n=27) were measured and those exhibiting an MFI-B greater than or equal to 10 times the median of any of the 10 CD antigens in either sample type were eliminated as outliers (n=7). The remaining 64 controls were used to construct cut off values (C_0_) to determine positivity for CD patient samples. Positivity was determined to be the average plus five standard deviations for MENSA and four standard deviations for serum. CDI samples were selected within 12 DPV and tested against the 10 CD antigens. If a patient had more than one blood draw during the analysis time period, the sample exhibiting the greatest sum of 9 CD antigens was selected (TcdA-CROP, TcdA-GTD, TcdB-CROP, TcdB-GTD, CDTa, CDTb, GDH, Cwp84, FLiC). TcdBvir-CROP was excluded from this determination due to its similar values to TcdB-CROP. MENSA and serum were analyzed separately to give the highest sum for each sample type. Of the 167 CDI patients, 39 had different time points for their MENSA and serum values but all were within 12 DPV. The majority of patients (128/167) had the same time point for their MENSA and serum samples. MFI-B was compared among patient groups and controls. Further analysis and data representation were undertaken using Microsoft Excel, GraphPad Prism, and JMP statistical analysis packages.

### Study Approval

The Emory and Dekalb Institutional Review Boards approved all protocols and procedures. Written informed consent was obtained from each patient prior to inclusion in the study.

## AUTHOR CONTRIBUTIONS

FL and JD designed and supervised the study. NH, SN, SO, FL, and JD wrote the manuscript with support/input/feedback from all authors. NH, SN, GK, and SO contributed to sample preparation, carried out the experiments, and analyzed the data. CK, AB, and PR acquired the patient samples. YW confirmed CDI diagnosis in patient samples. HW provided statistical guidance. SL oversaw patient recruitment. LC reviewed initial study design and data. MK provided detailed editing and thoughtful review of the manuscript. All authors contributed to the manuscript revision, read, and approved the submitted version.

## ACKNOWLEDGEMENTS

This work was supported by the Centers for Disease Control and Prevention through Phase I and Phase II SBIR Contracts #200-215-88233 titled “Antibody Secreting Cells (ASC) in Human *Clostridium difficile* Infection, Colonization, and Recurrence”. Special thanks are due to the clinical coordinators who assisted in this project. Coordinators at Emory Hospitals were Ms. Hinel Patel, Ms. Aja Bowser and Ms. Maya Lindsay. Our thanks also to the clinical coordinators at Dekalb Hospital and IDS Atlanta, Mr. Travis Stewart and Ms. Julia Norton. Finally, a special thanks to Dr. Clifford McDonald at the Centers for Disease Control for his continued advice and support.

## Notes

**CONFLICTS OF INTEREST** Authors Natalie S. Haddad and Sophia Nozick are current full-time employees of MicroB-plex, Inc. Geena Kim and Shant Ohanian are former full-time employees of MicroB-plex, Inc., and generated samples and data reported in this manuscript. F. Eun-Hyung Lee, M.D., is the founder of MicroB-plex, Inc, and an Associate Professor at Emory University. She did not draw salary on this project, but she was actively engaged in grant-writing, project administration and production of the manuscript. John L. Daiss, Ph.D., works more than half-time for MicroB-plex, Inc. and also serves as a Research Associate Professor at the University of Rochester Medical Center. L. Edward Cannon was the CEO of MicroB-plex, Inc., throughout the time of this study. The remaining authors declare that the research was conducted in the absence of any commercial or financial relationships that could be construed as a potential conflict of interest.

